# Time-lapse imaging of bacterial morphology facilitates classification of antimicrobial agents

**DOI:** 10.1101/2021.09.08.459470

**Authors:** Xudong Ouyang, Jelmer Hoeksma, Tjalling K. Siersma, Leendert W. Hamoen, Jeroen den Hertog

## Abstract

Antimicrobial resistance is a major threat to human health. Basic knowledge of antimicrobial mechanism of action (MoA) is imperative for patient care and for identification of novel antimicrobials. However, the process of antimicrobial MoA identification is relatively laborious. Here, we developed a simple, quantitative time-lapse fluorescence imaging method, Dynamic Bacterial Morphology Imaging (DBMI), to facilitate this process. It uses a membrane dye and a nucleoid dye to track the morphological changes of single *Bacillus subtilis* cells in response to antimicrobials for up to 60 min. DBMI of bacterial cells facilitated assignment of the MoAs of 14 distinct, known antimicrobial compounds to the five main classes. Using this method, we found that the poorly studied antimicrobial, harzianic acid, a secondary metabolite that we purified from the fungal culture of *Oidiodendron flavum*, targets the cell envelope. We conclude that DBMI is a simple method, which facilitates rapid classification of the MoA of antimicrobials in functionally distinct classes.

## Introduction

As the number of untreatable infections caused by multidrug-resistant “superbugs” is increasing globally, antimicrobial resistance is becoming a major threat to human health (Hancock, 2005; Blair et al., 2015). New antimicrobials with distinct working mechanisms are required to combat bacteria that have become resistant to all known antimicrobials. Although thousands of small molecules have now been screened for antibacterial activity in countless screening programs, the yield of useful compounds from these is relatively low (Payne et al., 2007). One of the reasons is the lack of insight into antimicrobial mechanism of action (MoA), which makes it difficult to determine the value of a new compound. Therefore, understanding the MoA is fundamental for identification of novel antimicrobials.

Transcriptomic and proteomic technologies have been developed to assess the MoA of antimicrobial agents in recent years (Van Duy et al., 2007; Silver, 2011). Transcriptomic analyses generate information about the expression of genes in response to antimicrobials (Pulido et al., 2016) and provide transcriptome-wide overviews of gene expression and are therefore used at the initial stages of target identification. Proteomic approaches generate overviews of protein levels, and changes in response to antimicrobial treatment may provide insight into the working mechanism (Brötz-Oesterhelt et al., 2005; Armengaud, 2013). Whereas these methods provide valuable information about antimicrobial targets, they do not necessarily provide information about the direct target(s) of antimicrobials. Because the time spent and cost involved in the genetic and omics-approaches is relatively high, they are not the best approaches for initial stages of target identification. A simple, cost-effective method would be preferred as a first approach.

Recently, imaging-based approaches started to be applied for antimicrobial MoA prediction. Bacterial cytological profiling was developed for Gram-negative bacteria to classify antimicrobial agents, utilizing high resolution fluorescent microscopy of cells stained with fluorescent dyes, or expressing fluorescent reporters (Nonejuie et al., 2013; Sun et al., 2018). For Gram-positive bacteria, a panel of GFP reporters was developed, covering many pathways and this panel was used successfully for in-depth analysis of antimicrobial targets (Müller et al., 2016; Wenzel et al., 2018; Zhu et al., 2018). Here, we developed a method to rapidly distinguish the effect of anti-Gram-positive bacterial compounds from all of the five main target classes, cell membrane, cell wall, protein, DNA or RNA. To achieve this, we used time-lapse imaging of fluorescent dye-stained *B. subtilis* to record dynamic changes. We improved the imaging protocol to make it simple and functional for bacterial long-term imaging. Using this method, dubbed Dynamic Bacterial Morphology Imaging (DBMI), we observed bacteria over time and established fluorescence intensities qualitatively and quantitatively. It allowed to rapidly distinguish between antimicrobials from all of the five different classes. Finally, we purified an antimicrobial activity from the fungal culture of *Oidiodendron flavum* and identified it to be harzianic acid, an understudied fungal secondary metabolite. Using DBMI, we established that harzianic acid targets the cell membrane and cell wall.

## Materials and Methods

### Strains and reagents

*B. subtilis* 168 was used for imaging in this study (Kunst et al., 1997). *O. flavum* (CBS 366.71) was obtained from Westerdijk Fungal Biodiversity Institute (the Netherlands) and used for biologically active compound production. Commercial antimicrobials were purchased from Sigma Aldrich (Table 1). FM4-64, 4′,6-diamidino-2-phenylindole (DAPI), SYTO-9 and SYTOX-Green were purchased from Thermo Fisher Scientific.

**Table 1.**
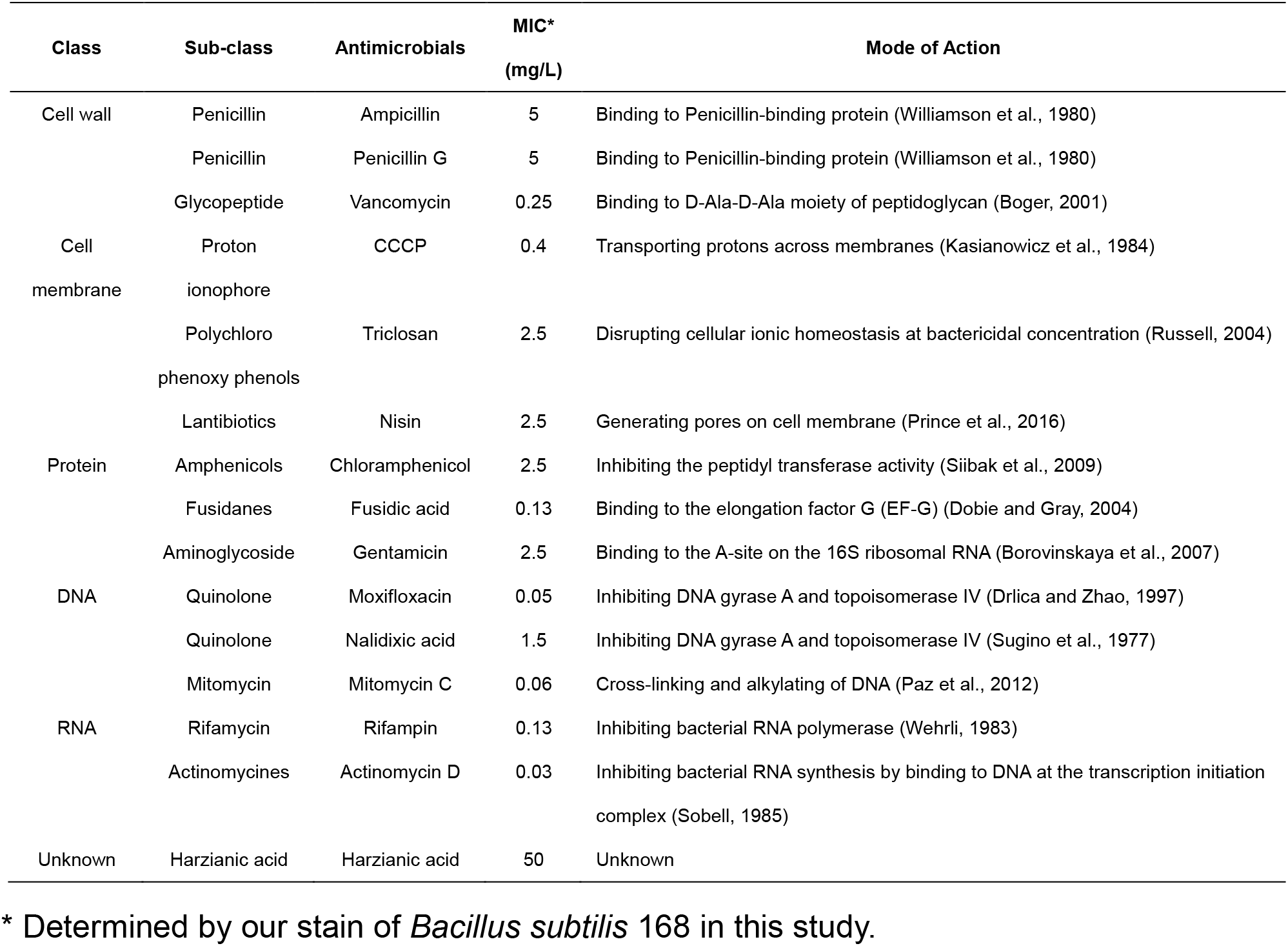
List of antimicrobials used in this study.

### Microdilution assay

MIC (Minimum Inhibitory Concentration) was determined by broth microdilution assay. Early exponential-phase cultures of *B. subtilis* were diluted 1:100 into Luria-Bertani (LB) medium, and distributed in 96-wells plates. Antimicrobials were tested 1:10 of the stock, then serially diluted with a factor 2. Bacterial growth was visually inspected after overnight incubation at 37 °C.

### Confocal microscopy

Microscopy was performed as described before with minor modification (Müller et al., 2016). Briefly, bacterial cultures in early exponential-phase were treated with antimicrobials (2.5 × MIC) or 1% dimethyl sulfoxide (DMSO) as control for up to 60 min. Afterwards, 1.5 µM FM4-64 was used for membrane staining. 5 µM DAPI, 50 nM SYTO-9 or 0.5 µM SYTOX-Green was applied to stain nucleoids. Samples were then immobilized on microscope slides covered with an agarose pad containing 1% agarose and LB medium, and imaged. Confocal microscopy was carried out using a Perkin Elmer UltraView VoX spinning disk microscope system and Volocity v6.3 software. Z-stack images were collected over a length of 3 µm with 0.2 µm intervals to acquire signals from the whole cells. Three independent experiments were done using each antimicrobial (biological triplicates). Images were analyzed using Fiji (Schindelin et al., 2012).

### Time-lapse microscopy

A single well agarose pad (2 mL in size) containing 0.75% agarose, 1.5 µM FM4-64, 50 nM SYTO-9 and LB medium was made by a trimmed syringe to immobilize bacterial cells on its surface. Bacterial culture in early exponential-phase was treated with 1.5 µM FM4-64 and 25 nM SYTO-9 for 10 min, and then 5 µL of it was transferred onto the agarose pad for imaging. Time-lapse images were collected using the spinning disk microscope system described above for 30 to 60 min with 3 min intervals. 400 µL LB medium with antimicrobials (2.5 × MIC) or 1% DMSO (control) was added into the well on the agarose pad after the second image was taken to make sure the cells were growing normally. Images were analyzed using Fiji.

### DBMI patterns

Stack images were merged by Z projection using max intensity and cell morphological patterns were measured by Fiji. A wide line (width: 20-pixel, i.e. 1.3 µm) was drawn to cover a whole cell and the intensity of both membrane signal and nucleoid signal over the line was acquired with the tool Plot Profile. The graphs of Plot Profile from each series of time-lapse imaging were then re-plotted into one heatmap with GraphPad Prism v8.4.1 to generate the DBMI patterns.

### Analysis of DBMI images

Stack images were merged by Z projection using max intensity. One cell unit was defined as a cell without any septa shown in the image of the initial time point. Cell status analysis was done by counting cells by eye in each of the three cell type categories (intact cell, no-nucleoid cell and disintegrated cell). Cell morphology analysis was done by measuring the length and intensity of both membrane and nucleoid using Fiji. Bar graphs and line chart were plotted using GraphPad Prism v8.4.1.

### Identification of biologically active compound

*O. flavum* was cultured on Malt Extract Agar (MEA) for 14 days. Secondary metabolites were extracted using ethyl acetate and separated using a Shimadzu preparative high performance liquid chromatography (HPLC) system with a C18 reversed phase Reprosil column (10 μm, 120 Å, 250 × 22 mm). The mobile phase was 0.1% trifluoroacetic acid in water (buffer A) and 0.1% trifluoroacetic acid in acetonitrile (buffer B). A linear gradient was applied of buffer B (5–95%) for 40 minutes. Fractions were collected and tested on *B. subtilis.* The active fraction was assessed for its purity through Shimadzu LC-2030 analytical HPLC using a Shimadzu Shim-pack GISTC18-HP reversed phase column (3 μm, 4.6 × 100 mm). LC-MS was performed on a Shimadzu LC-system connected to a Bruker Daltonics µTOF-Q mass spectrometer. High resolution mass spectrometry (HRMS) was measured on an LCT instrument (Micromass Ltd, Manchester UK). Elemental composition analyses were performed by Mikroanalytisch Labor Kolbe (Oberhausen, Germany). Finally, the compound was dissolved in 400 µL CDCl_3_ + 0,03% TMS and analyzed by Nuclear Magnetic Resonance (NMR) spectroscopy. More specifically, ^1^H-NMR, Hetronuclear Single Quantum Coherence (HSQC), Hetronuclear Multile-Bond Correlation (HMBC) and Correlation spectroscopy (COSY) spectra were measured at 600 MHz using a Bruker instrument. ^13^C-NMR was measured on the same instrument at 150 MHz.

## Results

### Static morphological differences did not fully distinguish the effect of antimicrobials on bacteria

Most antimicrobials for Gram-positives fall into one of five main classes based on their MoA: cell membrane, cell wall, protein, DNA or RNA (Peach et al., 2013). We set out to use morphological differences to distinguish MoAs between different classes of antimicrobials against Gram-positive bacteria. To this end, we selected one antimicrobial from each class. *B. subtilis* was treated for 60 min with the antimicrobials using 2.5 × MIC (Table 1) to ensure the inhibitory effects. The cells were stained with FM4-64 (red, cell membrane) and DAPI (blue, nucleoid), immobilized and imaged. Ampicillin (cell wall class) treatment for 60 min did not alter the appearance of cells compared to control treatment (Fig. 1). The other four antimicrobials induced variations both with respect to membrane and nucleoid staining. On the membrane, bright fluorescent foci appeared in response to all four antimicrobials, suggesting that the cytoskeleton was affected (Strahl et al., 2014). However, similar foci were also sometimes observed in the control, precluding the presence of foci as a criterion to distinguish between antimicrobial classes. Nucleoid staining was divided into two groups: decondensed nucleoids in response to CCCP (cell membrane class) and rifampin (RNA class), and shorter, more condensed nucleoids in response to moxifloxacin (DNA class) and chloramphenicol (protein class). In addition, the nucleoid dye SYTOX-Green was applied separately to check the permeability of cells. However, no cells were shown to have permeable membrane in response to CCCP or ampicillin (Fig. S1A). Taken together, analysis of membrane and nucleoid staining of immobilized cells after treatment with antimicrobials for 60 min was not sufficient to definitively distinguish between the five classes of antimicrobials. Nevertheless, some differences were observed and we hypothesized that the response over time might be distinct. To investigate the dynamics of the cells’ responses to antimicrobials, we set up time-lapse imaging of bacteria stained with fluorescent dyes.

**Fig. 1.**
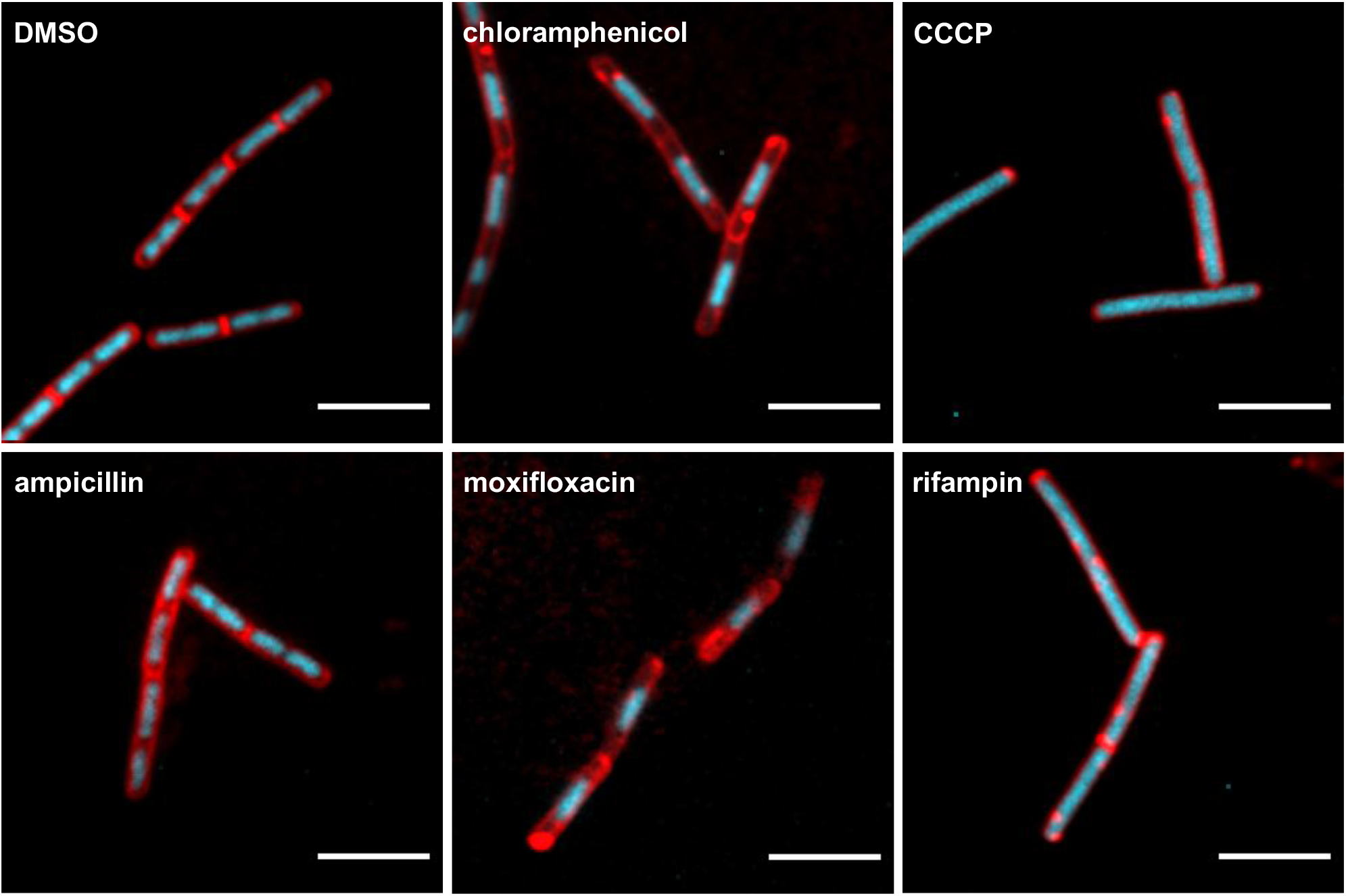
Cytological profiling of antimicrobial activities. *B. subtilis* cells were treated with an antimicrobial from one of the five main MoA classes or 1% DMSO (control) for 60 min. The antimicrobials we selected were: ampicillin for cell wall, CCCP for cell membrane, chloramphenicol for protein, moxifloxacin for DNA and rifampin for RNA. The concentrations of antimicrobials were 2.5 × MIC. Cells were stained with FM4-64 (red, cell membrane) and DAPI (blue, nucleoid), immobilized and imaged by confocal fluorescence microscopy. Representative images are shown. Scale bar is 5 µm.

### DBMI distinguished antimicrobials from distinct MoA classes

To acquire dynamic morphological patterns, SYTO-9 was used as nucleoid dye. We also designed the following workflow to enable time-lapse imaging of bacteria(Fig. 2). Images were taken every three minutes to prevent cells from laser damage, and two images were acquired before antimicrobials were added to confirm that the bacteria were growing healthily. To facilitate quantitative dynamic imaging of individual cells, the fluorescence intensity was integrated and quantified along the length of the cell and plotted. Control cells treated with DMSO showed steady growth over the course of 60 min (Fig. 3). DBMI was initially performed with the same antimicrobials as in the static bacterial imaging experiment. Images were collected for 60 min and resulting movies are shown in Movie S1-S6.

**Fig. 2.**
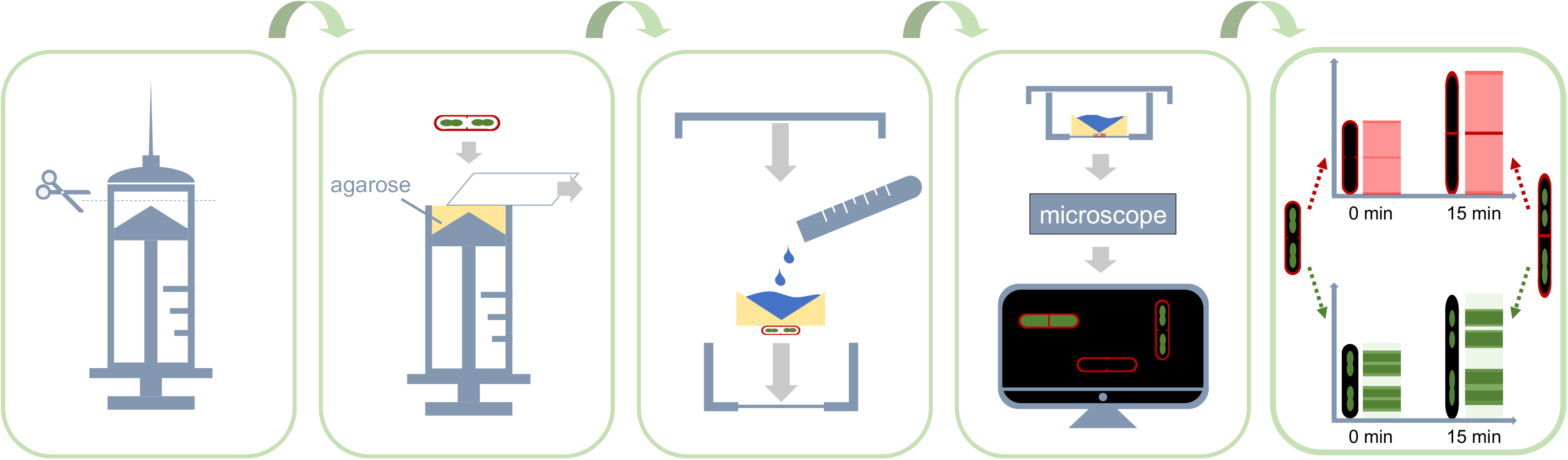
Workflow for time-lapse imaging of bacteria. A single well LB agarose pad was made using a trimmed syringe to immobilize bacterial cells on its surface. A glass slide was used to keep the agarose surface flat and smooth. Upon solidification of the agarose, the glass slide was removed and drops with bacterial cultures were added onto the surface. Once the drops were dry, the agarose pad with bacteria was transferred into a confocal dish for imaging. The thick agarose pad provided enough nutrition to sustain growing cells, allowing time-lapse imaging for hours, even overnight. The well on top of the agarose pad was used for easy addition of antimicrobial solutions to bacteria at any stage of the imaging process. After imaging, raw micrographs were converted into dynamic patterns by quantifying the fluorescence intensities of the membrane and nucleoid staining separately along the length of the bacteria.

**Fig. 3.**
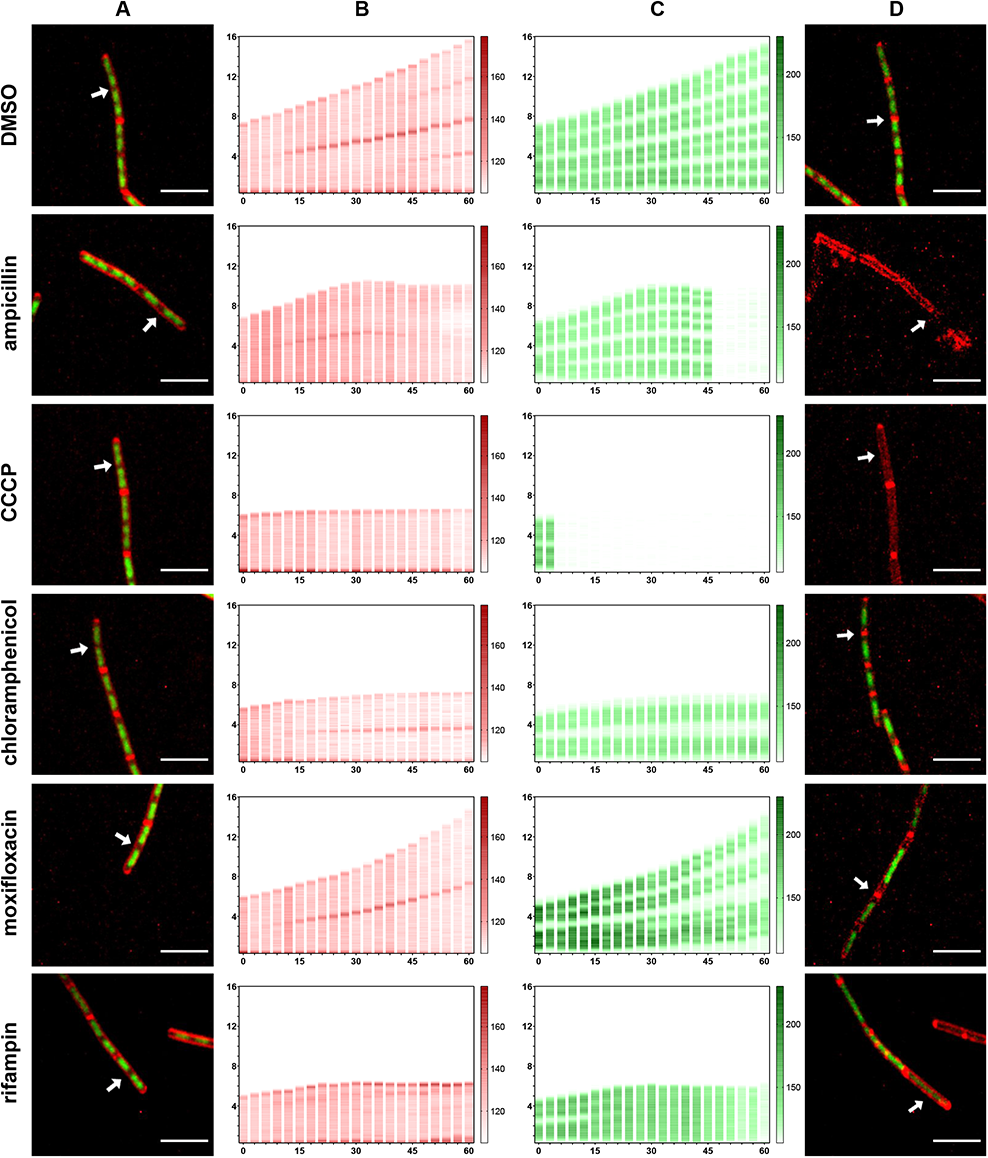
Distinct DBMI patterns of cells upon antimicrobial treatments. *B. subtilis* cells were treated with antimicrobials (2.5 × MIC) as indicated or 1% DMSO (control). The antimicrobials we selected were: ampicillin for cell wall, CCCP for cell membrane, chloramphenicol for protein, moxifloxacin for DNA and rifampin for RNA. Cells were stained with FM4-64 (red, cell membrane) and SYTO-9 (green, nucleoid) and imaged by time lapse confocal fluorescence microscopy for 60 min with 3 min intervals. Column A and D show the actual micrographs of the first and the last picture of the series (contrast was adjusted for individual graph to show details). The FM4-64 (column B) and SYTO-9 (column C) signals in a single cell (arrow in micrographs) were quantified over the length of the cell and plotted (y-axis, numbers in µm) over time (x-axis, numbers in min). Representative cells are shown. Scale bar is 5 µm.

Cells treated with different classes of antimicrobials had notably distinct profiles. The ampicillin and CCCP profiles stood out, because the nucleoid fluorescence signals suddenly became almost undetectable (Fig. 3). Ampicillin treatment inhibited cell growth and subsequently, the nucleoid fluorescence was lost. Strikingly, the membrane fluorescence at the septum started to decrease a few minutes after cell growth was inhibited. Part of the membrane fluorescence was lost at the end of the imaging series, showing that cells treated with ampicillin disintegrated, which was also evident from the actual micrograph (Fig. 3D). In response to CCCP, the membrane pattern remained normal. However, the nucleoid fluorescence intensity decreased sharply already after 6 min of treatment and cells stopped growing at the same time. The apparent loss of SYTO-9 staining was unexpected, because CCCP-treatment did not significantly affect DAPI-stained nucleoids (Fig. 1). Double labeling of nucleoids with SYTO-9 and DAPI showed that CCCP-treatment for 3 min already greatly reduced the SYTO-9 signal. In high contrast, SYTO-9 was still detectable. Yet, DAPI labeling appeared not to be affected in the same CCCP-treated cells (Fig. 4). Similar results were obtained after CCCP-treatment for 60 min (Fig. S1B). These results indicate that rapid reduction in SYTO-9 labeling of nucleoids was not due to loss of nucleoids from the cells, but rather to loss of SYTO-9 fluorescence.

**Fig. 4.**
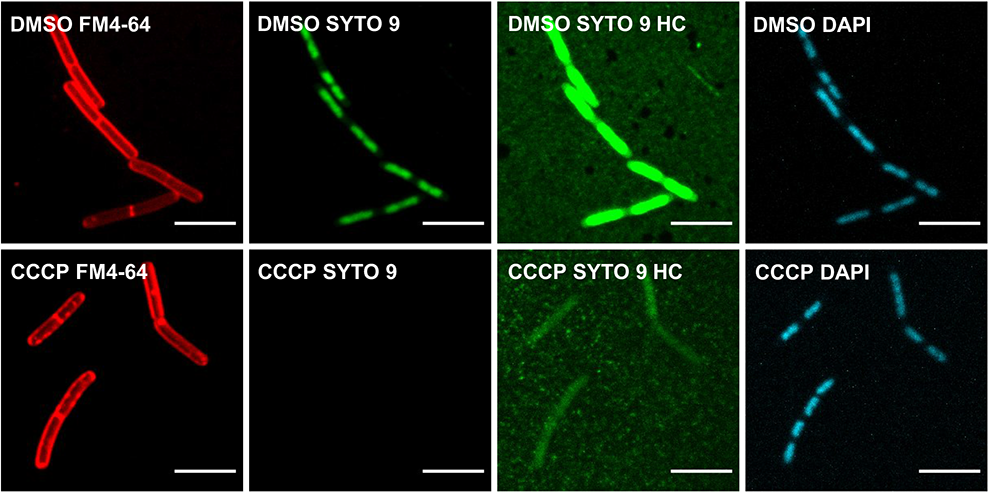
Dual nucleoid staining of cells following CCCP treatment for 3 min. *B. subtilis* cells were treated with CCCP (2.5 × MIC) or 1% DMSO (control) for 3 min. Cells were stained with FM4-64 (red, cell membrane), SYTO-9 (green, nucleoid) and DAPI (blue, nucleoid), immobilized and imaged by confocal fluorescence microscopy. Representative images are shown. Scale bar is 5 µm. SYTO-9 graphs were shown both in normal contrast and high contrast (HC).

Chloramphenicol and moxifloxacin induced shorter nucleoids relative to the cell size (Fig. 3), but the dynamics were different. In chloramphenicol treated cells, the extension of nucleoids was totally arrested, and the nucleoids compacted, resulting in an apparent increase in space between the nucleoids. In moxifloxacin-treated cells, the nucleoids still extended, but slower than the cells themselves, resulting in an increase in the space between nucleoids. Rifampin induced the nucleoid signal to gradually spread over the cells, suggesting that the nucleoids decondensed, which is in line with previous studies (Chen et al., 1996; Bakshi et al., 2015). Taken together, these results suggest that DBMI distinguished the MoA of antimicrobials from five major classes.

### Rapid loss of nucleoid staining was distinct in cells treated with antimicrobials from cell membrane and cell wall classes

In order to determine whether the DBMI profiles were conserved in each class, we tested 14 distinct antimicrobials from the five main antimicrobial classes (Table 1) using DBMI. All the profiles were first analyzed by eye for nucleoid visibility and membrane integrity to derive cell status, which was featured into three types: intact cells, having both visible membrane and nucleoid fluorescence; no-nucleoid cells, intact cells without visible nucleoid fluorescence; and disintegrated cells, no detectable nucleoid fluorescence and a disintegrated membrane. Cell status was evaluated over time to determine the changes in the ratios of each cell type during treatment.

As expected, antimicrobials within the cell membrane class (triclosan, nisin and CCCP) and the cell wall class (ampicillin, penicillin G and vancomycin) showed distinct profiles by cell status analysis and were readily distinguished from the other three classes (Fig. 5A, S1C). The percentage of intact cells decreased strongly after treatment with antimicrobials from the cell membrane and cell wall classes, compared to the other three classes. However, some antimicrobials from RNA and protein classes, including rifampin, gentamicin and chloramphenicol, also induced loss of nucleoid fluorescence in a small proportion of the cells (Fig. S1C). Quantification of nucleoid intensity of individual cells over time indicated that rifampin, gentamicin and chloramphenicol treatment resulted in gradual loss of nucleoid intensity (Fig. S1D). In contrast, treatment with cell membrane-active and cell wall-active antimicrobials resulted in a sharp decrease of nucleoid fluorescence intensity in each individual cell (Fig. 5B). Next, to distinguish the classes further, quantitative analysis of the changes in the membrane fluorescence intensity, cell length and the nucleoid length was done over time. Differences in membrane appearance and nucleoid shape were also recorded.

**Fig. 5.**
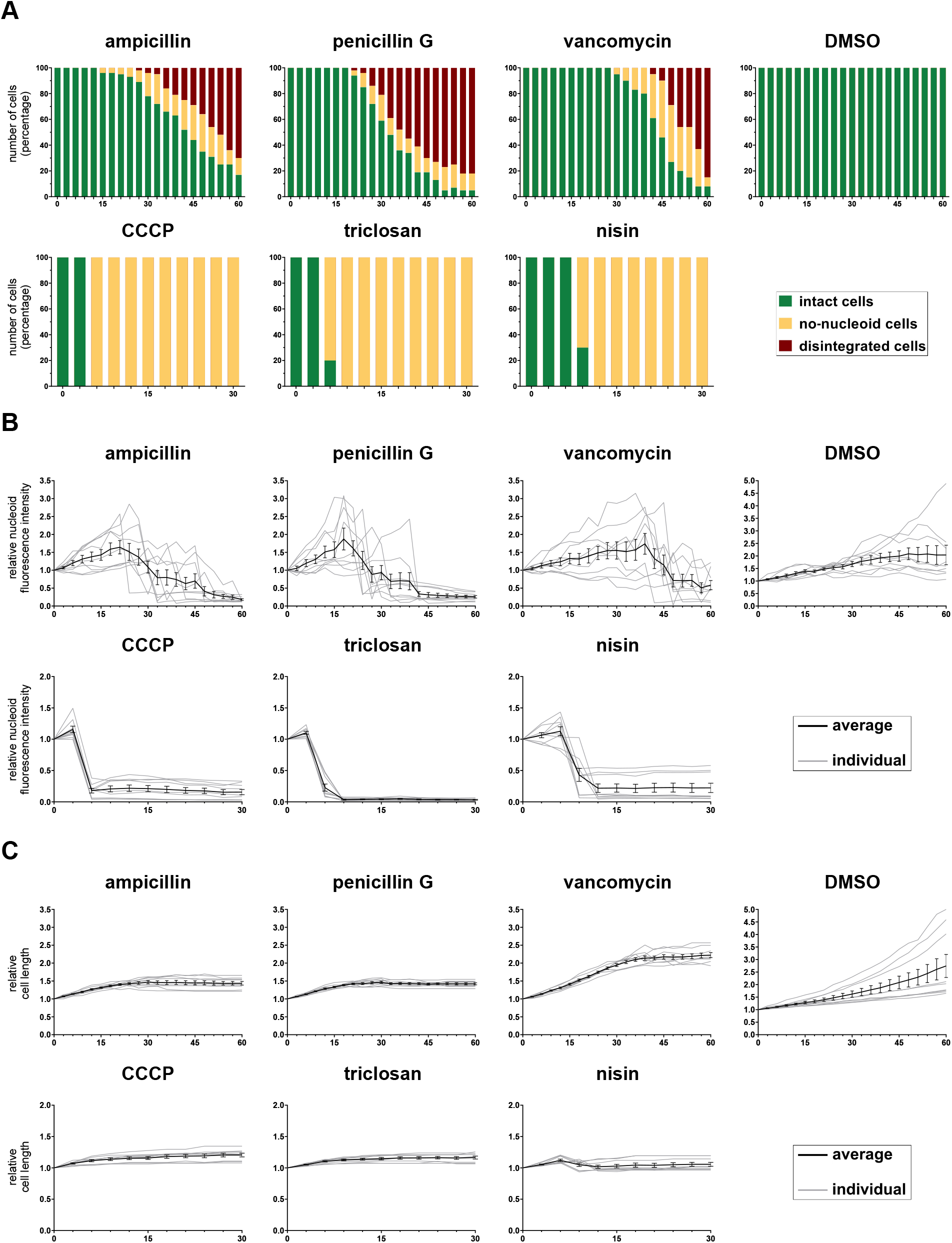
DBMI of cells treated with antimicrobials from the cell membrane and cell wall classes. *B. subtilis* cells were treated with antimicrobials (2.5 × MIC) as indicated or 1% DMSO (control). Cells were stained with FM4-64 (red, cell membrane) and SYTO-9 (green, nucleoid), and imaged by time lapse confocal fluorescence microscopy with 3 min intervals. Three distinct cell types were identified: intact cells (cells with visible membrane and nucleoid fluorescence), no-nucleoid cells (cells with apparently intact membrane, but without visible nucleoid fluorescence), and disintegrated cells (cells with disintegrated membrane and no detectable nucleoid). The number of cells of the three types were counted from biological triplicate imaging series (n > 40). In-depth cell morphology analysis was done on 9 cells in total per condition, i.e. 3 cells per biological triplicate. The cell length and nucleoid intensity data were quantified in these cells at each timepoint. (A) The number of cells in each cell type, intact cells (green), no-nucleoid cells (yellow) and disintegrated cells (red), was determined and plotted as percentage (y-axis, numbers in %) over time (x-axis, min). (B) Overall nucleoid intensity, i.e. the SYTO-9 fluorescence intensity inside a whole cell, corrected for background, was determined for individual cells over time and was depicted as percentage of the value at the start (t = 0 min) (y-axis) over time (x-axis, min). The mean of antimicrobial-treated cells was plotted in black with error bars representing the SEM. Gray lines represent individual antimicrobial-treated cells on which the mean was based. (C) Cell length was determined using the membrane signal (FM4-64) and was represented as percentage, relative to the initial length (y-axis, numbers in %) over time (x-axis, min). The mean of antimicrobial-treated cells was plotted in black with error bars representing the SEM. Gray lines represent individual antimicrobial-treated cells on which the mean was based.

To distinguish the cell membrane class from the cell wall class, we evaluated the percentage of intact cells (Fig. 5A) and the inhibition of cell growth (Fig. 5C). Cell wall-active antimicrobial treatment induced loss of nucleoid fluorescence and inhibition of cell growth after 15 - 30 min of treatment. Cell wall-active antimicrobials also induced disintegration of cells within 12 min after losing nucleoid fluorescence. Disintegrated cells were only observed upon treatment with cell wall-active antimicrobials, making this both a unique and a conserved trait for this class. In contrast, cell membrane-active antimicrobials induced simultaneous arrest of cell growth and loss of nucleoid fluorescence at an early stage of the antimicrobial treatment. These results indicate that the loss of nucleoid fluorescence is the first effect of cell membrane-active antimicrobials and distinguishes the class of cell membrane-active antimicrobials from the other classes. Like for CCCP, co-staining of nucleoids using SYTO-9 and DAPI indicated that loss of SYTO-9 fluorescence was not caused by loss of nucleoids themselves (Fig. S1E). Taken together, DBMI facilitates the distinction between cell wall and cell membrane active antimicrobials.

Next, we assessed differences between the cells treated with the three non-envelope-associated classes of antimicrobials. Since most of the cells remained intact during treatment (Fig. S1C), we investigated the potential differences in cell morphology, first focusing on changes in nucleoid length (Fig. 6A, S1F) in comparison with changes in cell length (Fig. 5C, 6B). To visualize this comparison, the ratio between nucleoid length and cell length was calculated (Fig. S1G). If not detectable, the nucleoid length was recorded as 0 µm. In response to cell wall and cell membrane active antimicrobials, nucleoid fluorescence was lost abruptly. In contrast, the two antimicrobials from the RNA class (rifampin and actinomycin D) induced an increase in nucleoid length compared to cell length, albeit with different dynamics. This indicated that an increased ratio of nucleoid length to cell length was conserved following treatment with antimicrobials from the RNA class, which is consistent with a previous report about nucleoid compaction (Cabrera et al., 2009). In comparison, treatment with antimicrobials from both the DNA class (moxifloxacin, mitomycin C and nalidixic acid) and the protein class (chloramphenicol, fusidic acid and gentamicin) reduced nucleoid length compared to cell length (Fig. S1G). For cells treated with antimicrobials from the protein class, nucleoid elongation stopped immediately, suggesting DNA elongation was blocked shortly after the antimicrobial was added. Since cell growth was not totally inhibited (fusidic acid) or was inhibited more slowly than the inhibition of nucleoid growth (chloramphenicol and gentamicin), the ratio of nucleoid length to cell length was reduced. The antimicrobials from the DNA class did not inhibit nucleoid elongation completely, but reduced it to some extent. Additionally, during treatment, the separation of nucleoids seemed to be poorly controlled and unusual shapes of nucleoids were often observed (e.g. Fig. 7A at 30 min), in line with known effects of these antimicrobials on DNA replication (Sugino et al., 1977; Drlica and Zhao, 1997; Paz et al., 2012). Hence, each class of antimicrobials we tested generated a unique and reproducible DBMI profile.

**Fig. 6.**
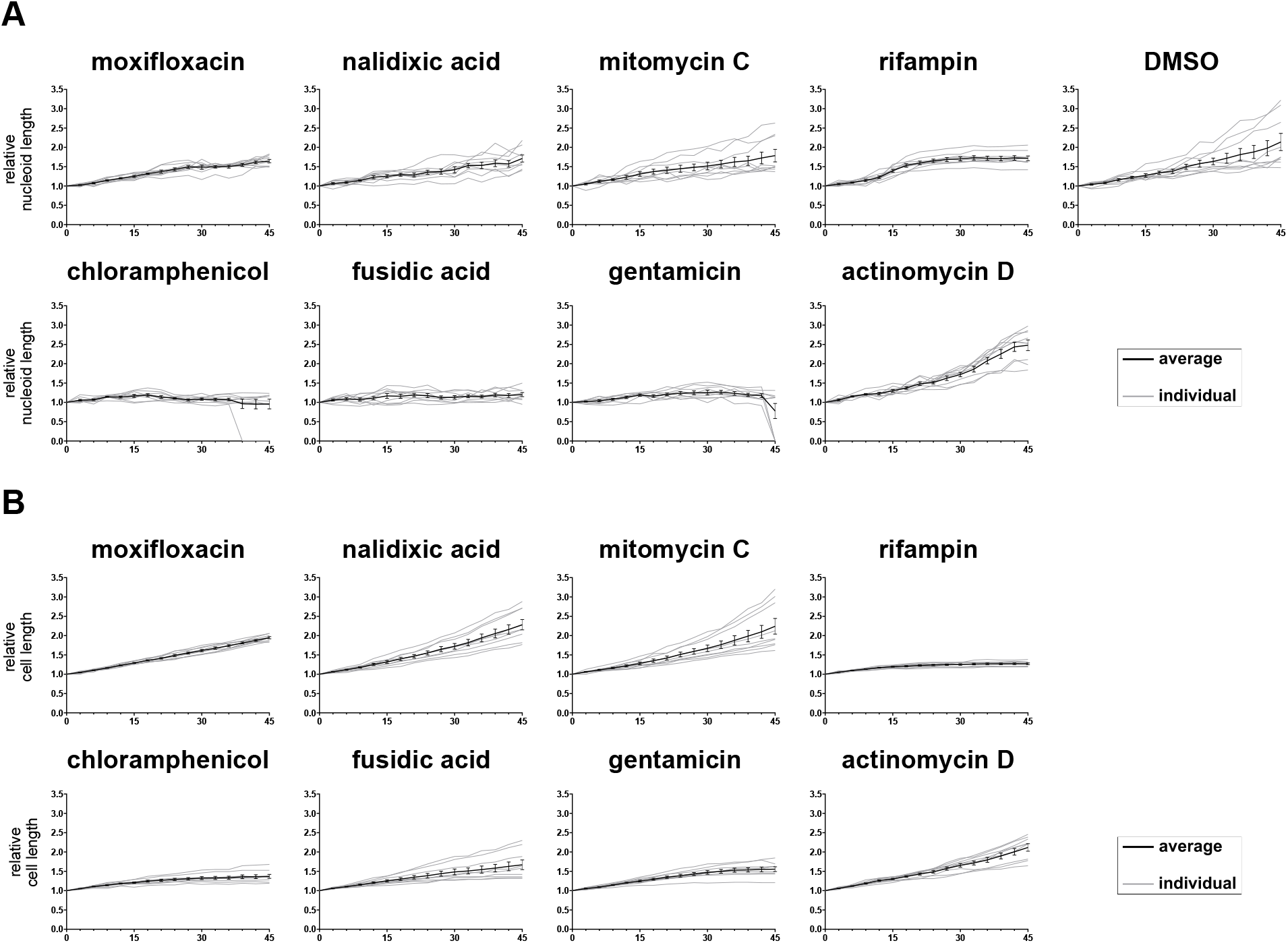
DBMI of cells treated with antimicrobials from the DNA, RNA and protein classes. *B. subtilis* cells were treated with antimicrobials (2.5 × MIC) as indicated or 1% DMSO (control). Cells were stained with FM4-64 (red, cell membrane) and SYTO-9 (green, nucleoid), and imaged by time lapse confocal fluorescence microscopy with 3 min intervals. Cell morphology data were analyzed from biological triplicate imaging series in triplicate, hence from 9 cells in total per antimicrobial and processed as described in the legend to Fig. 5. Nucleoid length, i.e. the length of all the nucleoids inside one cell added together, is shown in (A) and cell length in (B). In case nucleoid fluorescence was not detectable, the nucleoid length was recorded as 0 µm. The mean of antimicrobial-treated cells was plotted in black with error bars representing the SEM. Gray lines represent individual antimicrobial-treated cells on which the mean was based. DMSO control is depicted in Fig. 5C.

**Fig. 7.**
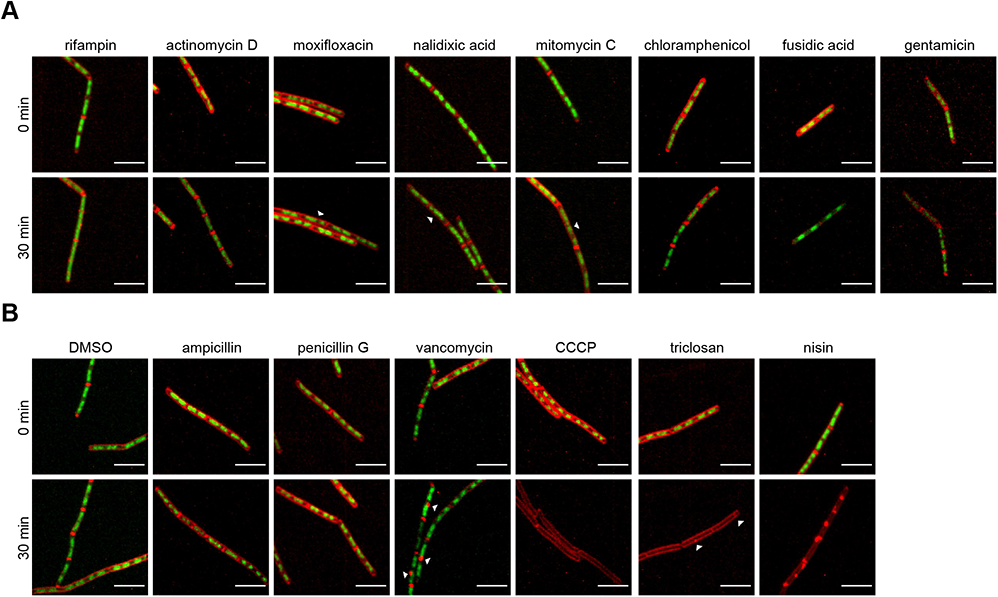
Cell membrane and nucleoid staining of cells after 0 min and 30 min treatment with different antimicrobials. Cells treated with antimicrobials from DNA, RNA and protein class are in (A) and from cell membrane, cell wall classes and DMSO control are in (B). Special patterns induced by moxifloxacin, nalidixic acid, mitomycin C, triclosan and vancomycin are indicated by arrow heads. Representative images are shown. Scale bar is 5 µm.

### DBMI distinguished sub-classes within the cell membrane, cell wall and RNA classes

Subsequently, we investigated whether DBMI was also able to distinguish sub-classes within the five main classes. In the cell membrane class, the growth curve of nisin-treated cells was notable, showing stabilized and even decreased cell size when the drug was added (Fig. 5C), which might due to loss of turgor pressure. In addition, big membrane blobs were observed after loss of nucleoid intensity at 30 min (Fig. 7B). Triclosan and CCCP generated highly similar profiles, but there was still a detectable morphological difference between the treated cells at 30 min (Fig. 7B). CCCP did not induce effects on the membrane, whereas triclosan induced small fluorescent membrane foci. These results are in line with their MoAs: nisin generates pores in the membrane, whereas triclosan and CCCP destroy membrane polarization, albeit at different levels (Kasianowicz et al., 1984; Russell, 2004; Prince et al., 2016). This indicates that DBMI may be used to distinguish between cell membrane-active antimicrobials.

When comparing the profiles within the cell wall class, vancomycin was found to generate a different profile than ampicillin and penicillin G. The cell growth inhibition in response to vancomycin was delayed by 20 min compared to the other two (Fig. 5C). Before cell growth inhibition at 30 min, some fluorescent blobs were detected on the membrane, indicating membrane aggregates during vancomycin treatment, which is distinct from cells treated with ampicillin or penicillin G (Fig. 7B). Ampicillin and penicillin G generated similar DBMI profiles and they both inhibit cross linking of peptidoglycans (Williamson et al., 1980), whereas vancomycin affects lipid II synthesis (Boger, 2001). Thus, we conclude that DBMI is able to distinguish antimicrobials of the cell wall class.

Rifampin of the RNA class induced rapid elongation of the nucleoid whereas cell length was almost constant, resulting in an increase in nucleoid length: cell length ratio (Fig. 6A, B and S1G), indicating that the nucleoids in these cells were decondensed (Cabrera et al., 2009) and almost fully filled the cells (Fig. 7A). Although the increase in the ratio of nucleoid length to cell length was also observed in actinomycin D treated cells, it was much slower than in rifampin treated cells, suggesting that actinomycin D has a different target to block RNA synthesis, which is consistent with previous reports (Wehrli, 1983; Sobell, 1985).

Next, we explored differences of the profiles of antimicrobial treatments within the protein class and the DNA class. Whereas most of the dynamic changes were similar among the three selected antimicrobials of the protein class, changes in nucleoid intensity were distinct for each antimicrobial (Fig. S1D). None of the fusidic acid treated cells showed a reduction in nucleoid intensity, whereas half of the chloramphenicol treated cells and all of the gentamicin treated cells showed a decrease in nucleoid intensity. This might be related to their different MoAs (Dobie and Gray, 2004; Borovinskaya et al., 2007; Siibak et al., 2009), but the large variability in nucleoid intensities in chloramphenicol treated cells precluded nucleoid intensity as a good parameter to distinguish between antimicrobials within the protein category. For the DNA class of antimicrobials, the responses were too similar to distinguish between them.

Taken together, DBMI facilitated distinction between antimicrobials from the five main classes, and allowed to distinguish sub-classes within the cell membrane, cell wall and RNA classes, using the scheme, depicted in Fig. 8.

**Fig. 8.**
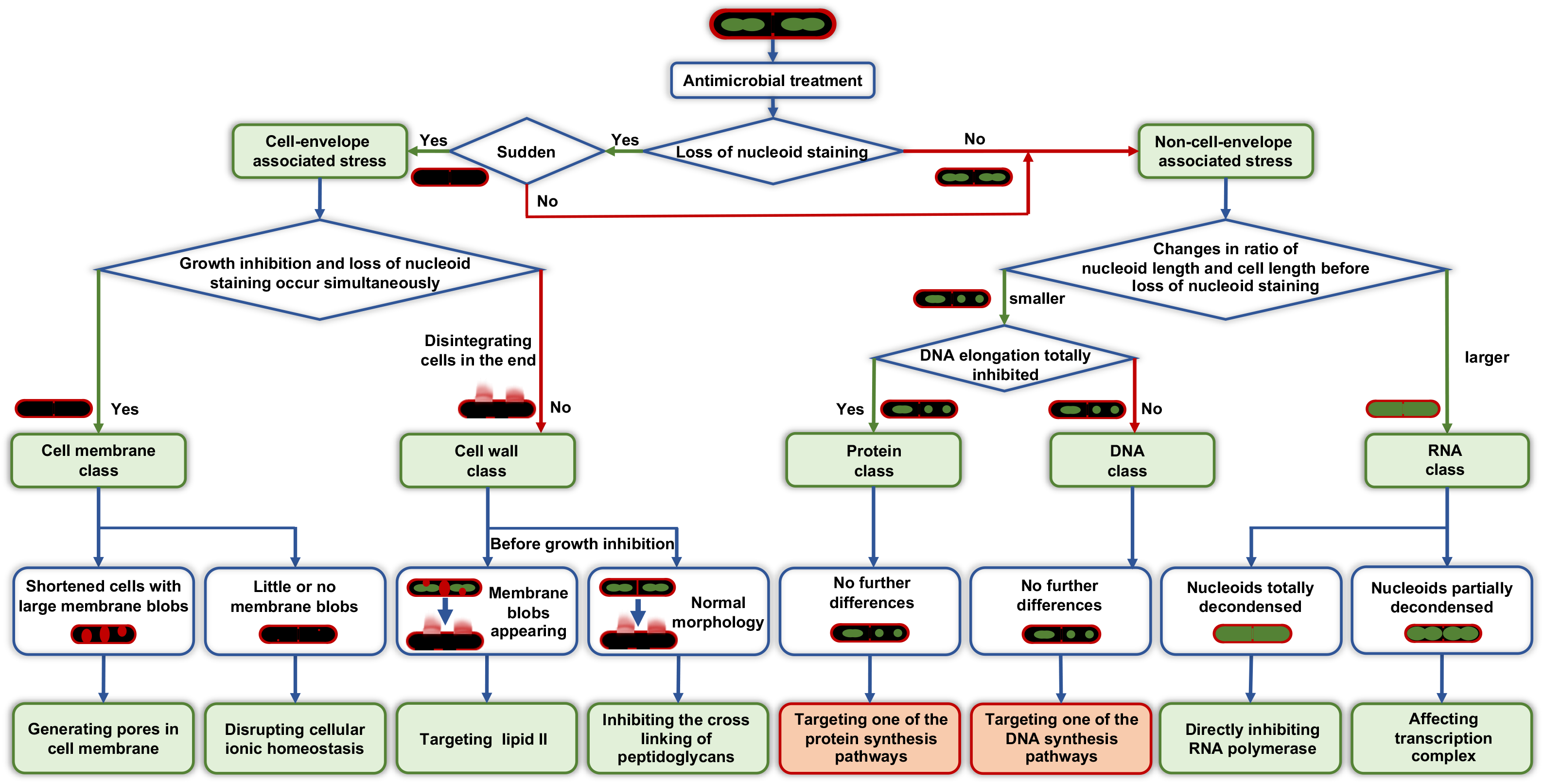
Schematic representation of the DBMI-guided antimicrobial MoA identification. Green boxes indicate well-identified categories or mechanisms, whereas red ones indicate not well-identified. DBMI distinguishes antimicrobial MoAs at three levels. Briefly, *B. subtilis* cells were treated with antimicrobials and imaged. At the first level, based on the loss of nucleoid staining, every antimicrobial was grouped into cell-envelope associated stress or non-cell-envelope associated stress. If the nucleoid staining was suddenly lost, they would fall into cell-envelope stress group. At the second level, five main MoA classes were identified. For the group of cell-envelope stress, if growth inhibition and loss of nucleoid staining occurred simultaneously, cell membrane was supposed to be affected. If growth inhibition occurred before loss of nucleoid staining, cell wall was determined to be affected if disintegrated cells were observed. For the group of non-cell-envelope associated stress, the changing of nucleoid length / cell length ratio before loss of nucleoid staining were measured. If the ratio was becoming larger during imaging, RNA synthesis was supposed to be affected. If the ratio was becoming smaller, the nucleoid growth had to be checked. If DNA elongation was inhibited, the antimicrobial would fall into protein class; if not, into DNA class. At the third level, sub-classes would be identified within cell membrane, cell wall and RNA classes. Protein and DNA classes could not be further distinguished because the differences within these classes were not evident. For the cell membrane class, if cells became shorter and large membrane aggregates were observed, probably pores were generated in the cell membrane. If there were little or no membrane blobs, the cellular ionic homeostasis was supposed to be disrupted. To further distinguish within cell wall class, we looked back to the time before growth inhibition. If cells were normal, the cross linking of peptidoglycans, which was the last step of cell wall synthesis, might be affected. If thicker membrane septa or big membrane blobs were observed, lipid II synthesis might be affected. For the RNA class, the extent of nucleoid decondensing was different between different targets. If RNA polymerase was targeted, nucleoids would be totally decondensed. If transcription complex was affected, the decondensing of nucleoids would be gentler.

### Identification of Harzianic Acid

Previously, we screened fungal metabolites from more than 10,000 fungi for bioactive compounds (Hoeksma et al., 2019). Here, we screened this library for antimicrobial activity against Gram-positive bacteria (*S. aureus*). One of the fungi, *O. flavum* (CBS 366.71), produced high antimicrobial activity. The active compound was purified and the chemical characteristics (Table 2, Fig. S2) were consistent with data previously reported for harzianic acid (Sawa et al., 1994; Vinale et al., 2014) (Fig. 9A).

**Fig. 9.**
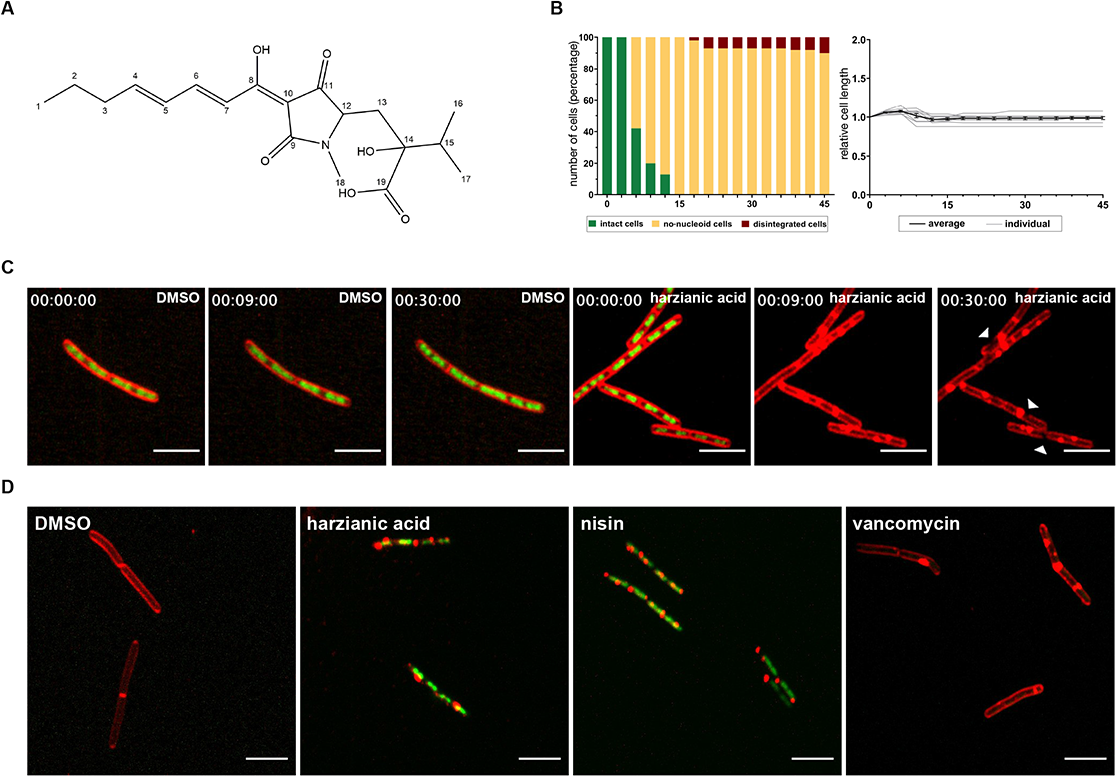
MoA identification of harzianic acid. (A) Chemical structure of harzianic acid. (B) Analysis of DBMI images of harzianic acid. *B. subtilis* cells were treated with harzianic acid (2.5 × MIC). Cells were stained, imaged, analyzed and plotted as in Fig. 5. (C) Examples of micrographs of cells at 0 min, 9 min and 30 min that were used for the quantifications in (B). Disintegrated cells at 30 min are indicated by arrow heads. DMSO treated cells are shown as control. Representative images are shown. Scale bar is 5 µm. (D) Cell permeability assay. *B. subtilis* cells were treated with antimicrobials (2.5 × MIC) or 1% DMSO (control) for 60 min. Cells were stained with FM4-64 (red, cell membrane) and SYTOX-Green (green, nucleoid), and imaged by confocal fluorescence microscopy. Representative images are shown. Scale bar is 5 µm.

**Table 2:** Assignments NMR-shifts, HMBC and COSY couplings for harzianic acid (in CDCl_3_)

| Harzianic acid (CDCl<sub>3</sub>) |  |  |  |  |
| --- | --- | --- | --- | --- |
| # | δC<sup>a</sup> | δH<sup>b</sup> | HMBC<sup>b</sup> | COSY<sup>b</sup> |
| 1 | 13.7 | 0.95 (t), 3H | 2, 3 | 2 |
| 2 | 21.8 | 1.50 (m), 2H | 1, 3, 4 | 1, 3 |
| 3 | 35.5 | 2.24 (dd), 2H | 1, 2, 4, 5 | 3, 4 or 5<sup>c</sup> |
| 4 | 149.9 | 6.37<sup>c</sup> (m), 1H | 2, 3, 5, 6, 7<sup>c</sup> | 3, 6<sup>c</sup> |
| 5 | 129.6 | 6.38<sup>c</sup> (m), 1H | 2, 3, 6, 7<sup>c</sup> | 3, 6<sup>c</sup> |
| 6 | 147.6 | 7.55 (m), 1H | 4, 5, 7, 8 | 4 or 5<sup>c</sup>, 7 |
| 7 | 119.1 | 7.00 (d, *J*=15.1 Hz), 1H | 5, 6, 8, 10 | 6 |
| 8 | 176.7 | - | - | - |
| 9 | 173.2 | - | - | - |
| 10 | 99.7 | - | - | - |
| 11 | 197.3 | - | - | - |
| 12 | 64.1 | 3.63 (dd, *J*= 10.6, 1.0 Hz), 2H | 9, 11, 13, 14, 18, | 13 |
| 13 | 33.8 | 1.89 (dd), 1H<br>2.48 (d), 1H | 11, 12, 14, 15, 19 | 12 |
| 14 | 79.9 | - | - | - |
| 15 | 36.0 | 2.02 (m), 1H | 13, 14, 16, 17, 19 | 16 or 17<sup>c</sup> |
| 16 | 17.5 | 0.99<sup>c</sup> (m), 3H | 14, 15<sup>c</sup> | 15 |
| 17 | 16.2 | 0.99<sup>c</sup> (m), 3H | 14, 15<sup>c</sup> | 15 |
| 18 | 26.6 | 2.97 (s), 3H | 9, 12 | - |
| 19 | 176.3 | - | - | - |

<sup>a</sup>= measured at 150 MHz, <sup>b</sup>= measured at 600 MHz, <sup>c</sup>= overlapping signals

Harzianic acid: C_19_H_27_NO_6_, dark yellow powder. HRMS: found 388.1750 (M+Na), calculated 388.1736 for C_19_H_27_NO_6_Na. Elemental composition analyses: C 61,4%; O 21,8%; H 7,0%; N 3,8%. NMR (600 MHz, CDCl_3_): See Table 2. UV-Vis λ_max_: 244 nm, 363 nm.

### MoA Identification of Harzianic Acid

Little is known about harzianic acid, a compound with antimicrobial activity that was isolated from fungi decades ago (Sawa et al., 1994). Harzianic acid also has antifungal activity and plant growth promoting activity (Vinale et al., 2009, 2014; De Filippis et al., 2020; De Tommaso et al., 2020). The antimicrobial MoA of harzianic acid is unknown. DBMI was used to get insight into the MoA of harzianic acid. Imaging data were analyzed to assess cell status and cell morphology (Fig. 9B, S1). Harzianic acid induced loss of nucleoid fluorescence at an early stage of treatment. Cell growth was totally inhibited at an early stage as well. This was reminiscent of the initial action of cell membrane-active antimicrobials. Cell length actually appeared to decrease when the drug was added, suggesting loss of turgor pressure. Big membrane aggregates were observed when intact cells lost their nucleoid fluorescence, i.e. from 9 min onwards (Movie S7). These effects were similar to the effects of nisin, suggesting that harzianic acid generated pores in the membrane. Cell status data also indicated that part of the cells disintegrated. Note the gaps along the membrane at 30 min (Fig. 9C) or from 18 min onwards (Movie S7). Whereas disintegrated membranes were evident in nisin-treated cells as previously shown in static bacterial imaging (Lamsa et al., 2012), we did not observe disintegrated cells in response to nisin by DBMI up to 30 min treatment. Disintegration of part of the cells in response to harzianic acid may therefore suggest that harzianic acid also partially affects the cell wall.

The final disintegrated cell membrane showed similarities to the cell membrane of vancomycin treated cells, but the dynamics suggested that pore formation in the membrane was the initial effect of harzianic acid. To investigate this, non-cell permeable SYTOX Green was used to stain nucleoids. Evidently, treatment with harzianic acid or nisin, but not vancomycin or vehicle control (DMSO) induced SYTOX Green staining of the nucleoid (Fig. 9D). In conclusion, harzianic acid generated pores in the membrane and eventually appeared to affect the cell wall.

## Discussion

Here we used time-lapse imaging of live, stained Gram-positive bacteria to rapidly distinguish the MoA classes of antimicrobial agents. In order to acquire dynamic profiles of bacteria under different conditions, we used SYTO-9 instead of DAPI, because DAPI is toxic to bacterial cells. SYTO-9 shows low bacterial toxicity, fast staining, high fluorescence intensity in the green spectrum and therefore good compatibility with FM4-64 fluorescence. We also developed an imaging protocol (Fig. 2), which is based on a previously described technique (de Jong et al., 2011). A thick agarose pad provides enough nutrition to sustain cell growth and the well on top of the agarose is convenient for adding antimicrobial solutions to bacteria at any stage of the imaging process. In addition, the preparation is fast because no fridge-cool-down or precision-work steps are required. Using this imaging technique, which we called DBMI, we are able to reliably predict the MoA of antimicrobials at three levels (Fig. 8).

At the first level, antimicrobials were divided into two groups, based on whether they exert their effect on the cell envelope or not. All of the inhibitors that induce cell envelope stress induced a sudden loss of SYTO-9 nucleoid fluorescence, which is distinct from the effects of antimicrobials with non-cell-envelope stress targets. Rapid loss of SYTO-9 staining might be caused by sudden changes in intracellular milieu due to a compromised cell envelope, which affects SYTO-9 fluorescence. Which environmental factor(s) affect SYTO-9 fluorescence, be it membrane potential, pH, ionic profile or osmolarity, remains to be determined (Auty et al., 2001; Stiefel et al., 2015).

At the next level, antimicrobials were further distinguished into five distinct classes. For antimicrobials from the cell membrane class, cell growth arrest and loss of nucleoid staining occurred simultaneously, which was not observed in cells treated with cell wall active antimicrobials. In addition, cell wall active antimicrobials led to disintegrated cells eventually, which is presumably due to the decrease in peptidoglycan synthesis and increase in autolysins (Kohanski et al., 2010). Interestingly, this pattern was not observed in static bacterial imaging, e.g. ampicillin treated cells in Fig. 1. This discrepancy is likely caused by the different imaging methods that were used. Time lapse imaging of the same cell showed disintegration of the cell (Fig. 3D), whereas treatment in liquid medium for 60 min and subsequent transfer for imaging may cause disintegrated cells to go unnoticed and only intact cells are imaged (Fig. 1). The non-cell-envelope associated antimicrobials from different classes did not induce sudden loss of nucleoid fluorescence. Cells treated with RNA synthesis inhibitors showed decondensed nucleoids, whereas antimicrobials affecting the protein or DNA class both induced compacted nucleoids, consistent with earlier reports (Bylund et al., 1993; Van Helvoort et al., 1996; Bakshi et al., 2015). Unusual nucleoid shapes and shorter nucleoids may be due to a DNA segregation block and/or dissected nucleoids by division (Georgopapadakou and Bertasso, 1991; Chen et al., 1996; Drlica et al., 2008).

At the third level, antimicrobials from cell membrane, cell wall and RNA classes were further distinguished into sub-classes. In the cell membrane class, the appearance of the membrane seemed to be related to the extent of the permeability. From CCCP to triclosan to nisin, the permeability increased and apparently, so did the extent of the membrane aggregation. Notable among them was triclosan, which has multiple targets in the cytoplasm and membrane. High concentrations of triclosan have bactericidal effects and are mediated by membrane targets (Russell, 2004). Here, we used high concentrations (2.5 × MIC), which had bactericidal effects. DBMI classified triclosan in the cell membrane class, which is consistent with bactericidal concentrations of triclosan acting on membrane targets. In the cell wall class, vancomycin binds to different target groups than the penicillins (Williamson et al., 1980; Boger, 2001), which is in line with the observed aggregation of membrane only in response to vancomycin in the cell wall class. In the RNA category, due to the differences in targets (Wehrli, 1983; Sobell, 1985), the extent of decondensing of nucleoids was shown to be different. Inhibiting RNA polymerase by rifampin might affect RNA synthesis faster than actinomycin D binding to DNA at the transcription initiation complex, resulting in totally decondensed nucleoids in rifampin treated cells and only partially decondensed nucleoids in actinomycin D treated cells.

## Conclusion

DBMI facilitates profiling of changes over time in cell morphology and viability. With the obtained parameters, antimicrobials may be classified at three levels. DBMI does not require prior knowledge of the antimicrobial MoAs. Hence, DBMI may be used to rapidly distinguish the MoA class and subclass of known and unknown antimicrobials.

## Acknowledgements

The authors would like to thank Anko de Graaff of the Hubrecht Imaging Centre for help with imaging, and Samantha van der Beek and Maja Solman for help with developing the imaging protocol. This project was supported by the Chinese Scholarship Council (CSC).

