## Supplemental Material for "Time-lapse imaging of bacterial morphology facilitates classification of antimicrobial agents"

### Legends to supplemental movies S1-S7

**Dynamic profiling patterns of cells upon antimicrobial treatment.** *B. subtilis* cells were stained with FM4-64 (red, cell membrane) and SYTO-9 (green, nucleoid) and imaged by time lapse confocal fluorescence microscopy for 60 min with 3 min intervals. Cells were treated with 1% DMSO (control, Movie S1) or antimicrobials (2.5 × MIC): ampicillin, Movie S2; CCCP, Movie S3; chloramphenicol, Movie S4; moxifloxacin, Movie S5; Rifampin, Movie S6; harzianic acid, Movie S7. Antimicrobials were added after 6 min; time is indicated in the top left corner as hh:mm:ss. Representative cells are shown.

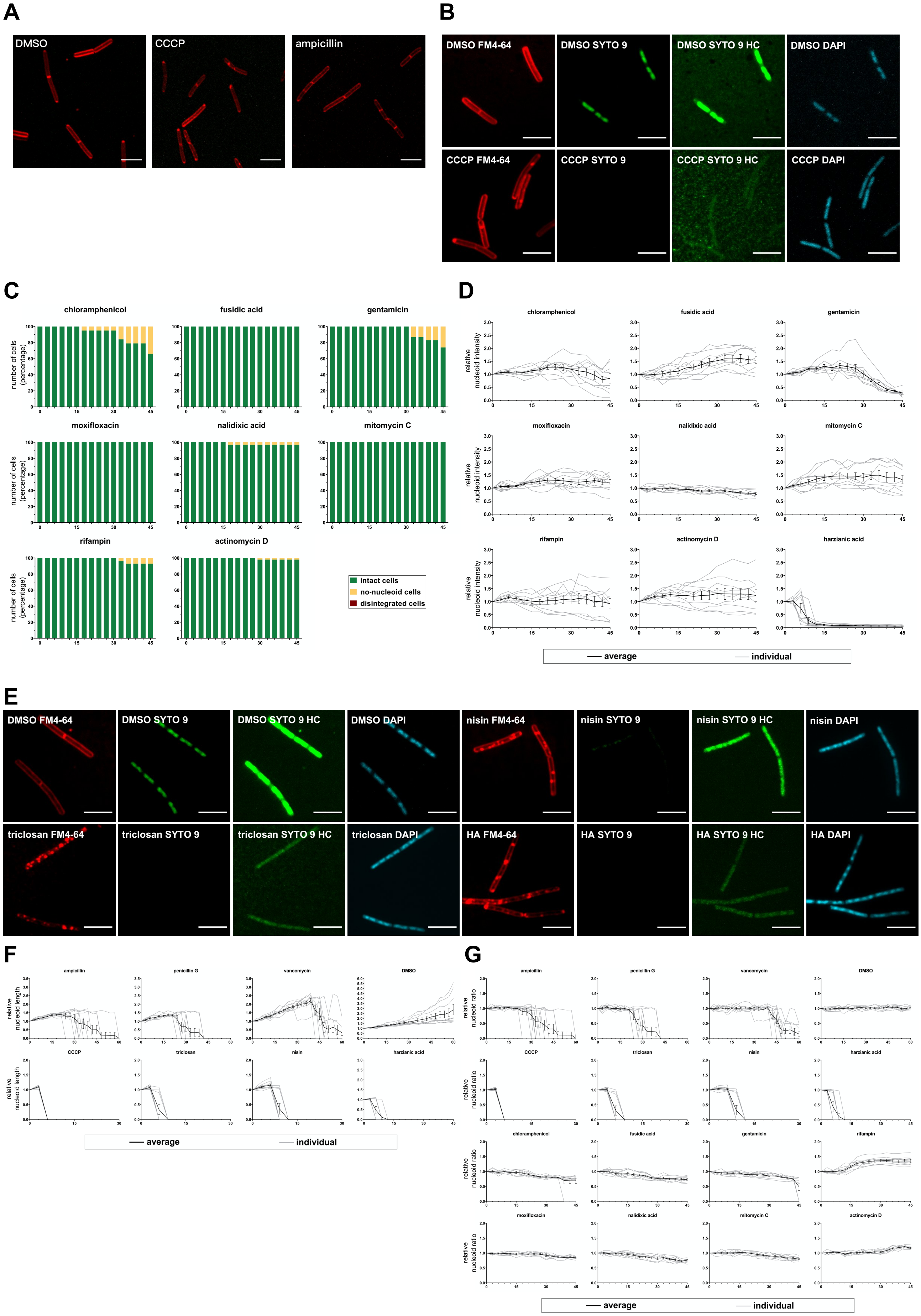
A
DMSO
CCCP
ampicillin
B
DMSO FM4-64
DMSO SYTO 9
DMSO SYTO 9 HC
DMSO DAPI
CCCP FM4-64
CCCP SYTO 9
CCCP SYTO 9 HC
CCCP DAPI
C
chloramphenicol
fusidic acid
gentamicin
moxifloxacin
nalidixic acid
mitomycin C
rifampin
actinomycin D
number of cells (percentage)
intact cells
no-nucleoid cells
disintegrated cells
D
harzianic acid
relative nucleoid intensity
average
individual
E
nisin FM4-64
nisin SYTO 9
nisin SYTO 9 HC
nisin DAPI
triclosan FM4-64
triclosan SYTO 9
triclosan SYTO 9 HC
triclosan DAPI
HA FM4-64
HA SYTO 9
HA SYTO 9 HC
HA DAPI
F
penicillin G
vancomycin
triclosan
nisin
relative nucleoid length
G
relative nucleoid ratio

**Fig. S1. Additional BCP and DBCP profiles.**

(A) Cell permeability assay. *B. subtilis* cells were treated with antimicrobials (2.5 × MIC) or 1% DMSO (control) for 60 min. Cells were stained with FM4-64 (red, cell membrane) and SYTOX-Green (green, nucleoid), and imaged by confocal fluorescence microscopy. Representative images are shown. Scale bar is 5 μm.

(B) Dual nucleoid staining of cells following CCCP treatment for 60 min. *B. subtilis* cells were treated with CCCP (2.5 × MIC) or 1% DMSO (control) for 60 min. Cells were stained with FM4-64 (red, cell membrane), SYTO-9 (green, nucleoid) and DAPI (blue, nucleoid), and imaged by confocal fluorescence microscopy. Representative images are shown. Scale bar is 5 μm. SYTO-9 graphs were shown both in normal contrast and high contrast (HC).

(C) Changes in cell status upon antimicrobial treatment. *B. subtilis* cells were treated with antimicrobials (2.5 × MIC) as indicated. Cells were stained with FM4-64 (red, cell membrane) and SYTO-9 (green, nucleoid), and imaged by time lapse confocal fluorescence microscopy with 3 min intervals. Three distinct cell types were identified: intact cells (cells with visible membrane and nucleoid fluorescence), no-nucleoid cells (cells with apparently intact membrane, but without visible nucleoid florescence) and disintegrated cells (cells with disintegrated membrane and no detectable nucleoid). The number of cells of the three types were counted from biological triplicate imaging series (n > 20). The ratio of cell types, intact cells (green), no-nucleoid cells (yellow) and disintegrated cells (red), was determined and plotted as percentage (y-axis, numbers in %) over time (x-axis, min). See Fig. 5A for DMSO control as this figure is an extension.

(D) Changes in nucleoid intensity upon antimicrobial treatment. *B. subtilis* cells were treated with antimicrobials (2.5 × MIC) as indicated. Cells were stained and imaged as in (C). Cell morphology data were collected from 9 cells in total per treatment, i.e. technical triplicates from biological triplicate imaging series. Overall nucleoid intensity, i.e. the SYTO-9 fluorescence intensity inside a whole cell, corrected for background, was determined for individual cells over time and was depicted as percentage of the value at the start (t = 0min) (y-axis) over time (x-axis, min). The mean of antimicrobial-treated cells was plotted in black with error bars representing the SEM. Gray lines represent individual antimicrobial-treated cells on which the mean was based. See Fig. 5B for DMSO control as this figure is an extension.

(E) Dual nucleoid staining of cells following treatment with antimicrobials from cell membrane class or harzianic acid (HA) for 3 min. *B. subtilis* cells were treated with antimicrobials (2.5 × MIC) as indicated or 1% DMSO (control) for 3 min. Cells were stained with FM4-64 (red, cell membrane), SYTO-9 (green, nucleoid) and DAPI (blue, nucleoid), and imaged by confocal fluorescence microscopy. Representative images are shown. Scale bar is 5 µm. SYTO-9 graphs were shown both in normal contrast and high contrast (HC).

(F) Changes in nucleoid length upon antimicrobial treatment. *B. subtilis* cells were treated with antimicrobials (2.5 × MIC) as indicated or 1% DMSO (control). Cells were stained, imaged and analyzed as in (D). Nucleoid length, the addition of the length of all the nucleoids inside one cell, was determined for individual cells and depicted as percentage of the value at the start (t = 0min) (y-axis) over time (x-axis, min). The mean of antimicrobial-treated cells was plotted in black with error bars representing the SEM. Gray lines represent individual antimicrobial-treated cells on which the mean was based. This figure is an extension of Fig. 6A, where the first 45 min of DMSO control has already shown.

(G) Changes in the ratio of nucleoid length to the cell length upon antimicrobial treatment. *B. subtilis* cells were treated with antimicrobials (2.5 × MIC) as indicated or 1% DMSO (control). Cells were stained, imaged and analyzed as in (D). The ratio of the nucleoid length to the cell length, was calculated for individual cells over time and was depicted as percentage of the value at the start (t = 0min) (y-axis) over time (x-axis, min). The mean of antimicrobial-treated cells was plotted in black with error bars representing the SEM. Gray lines represent individual antimicrobial-treated cells on which the mean was based.

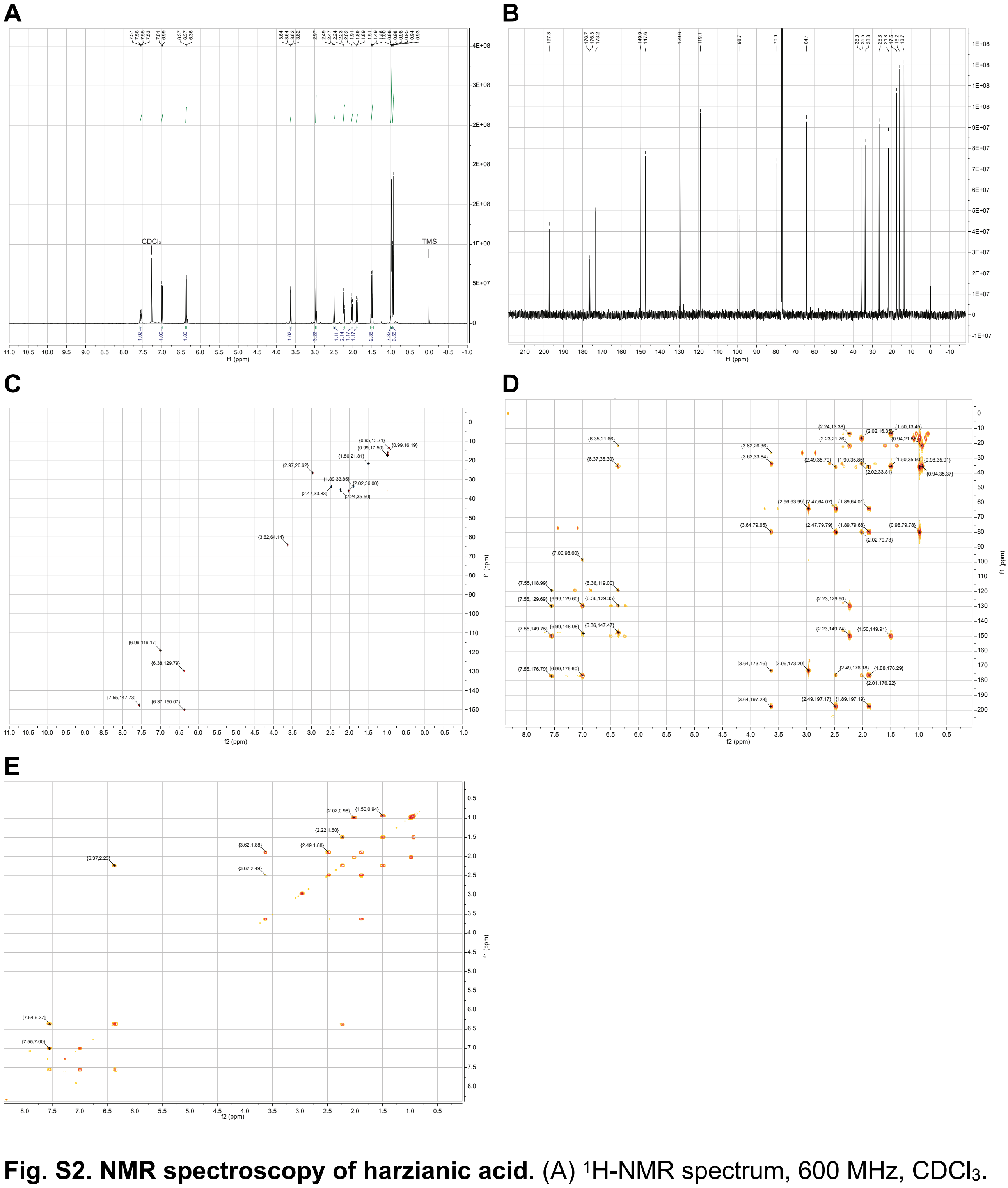


**Fig. S2. NMR spectroscopy of harzianic acid.** (A) $^{1}$H-NMR spectrum, 600 MHz, $CDCl_3$. (B) $^{13}$C-NMR spectrum, 150 MHz, $CDCl_3$. (C) HSQC spectrum, 600 MHz, $CDCl_3$. (D) HMBC spectrum, 600 MHz, $CDCl_3$. (E) COSY spectrum, 600 MHz, $CDCl_3$.
